# Cytotoxic Effect of Silver Ions in Combination with Peanut’s leaf extract on Cells

**DOI:** 10.1101/2021.07.24.453619

**Authors:** Yeonjo Jung

## Abstract

As modern people suffered to various infectious diseases such as MERS and Covid 19, people interest in nature-driven health care has increased. Accordingly, expectations for natural therapeutics that can have a beneficial effect on the human body without being processed have increased. Buckwheat, peanut, and pine are representatives. Among them, peanuts contain polyphenols and flavonoids which are used as nutritional foods in various forms. For example, resveratrol is activated by the CYP1B1 enzyme in peanut sprouts and converted to piceatannol that can act on anticancer activity. Accordingly, increasing the content of polyphenols in peanuts makes possible to produce useful substances that are harmless to the human body as well as the environment. Therefore, I conducted research to find an effective way to increase the content of polyphenols. This study projects an effective way to secure peanuts with high polyphenol content based on the characteristics that the content of polyphenols increases when plants are wounded. Comparative studies were conducted through wounding, and it was confirmed that the polyphenol content increased in the wounded plants. Subsequently, it was confirmed that the content of polyphenol increased through the DPPH experiment, and the content of flavonoids also increased. Using the data of this study can be used to produce peanuts with high polyphenol content. To find an effective method for cultivating peanuts containing high concentration of phenolic compounds, I compared two groups of plants with different conditions. First group was consisted of normal plants with no treatment and second group was consisted of wounded plants. I treated by using these conditions in each group for two days and the results showed that group 2 has high concentration of flavonoid and phenolic compounds.

To verify this result, I conducted DPPH assay which provides convincing evidence that group 2 has justifiable anti-inflammatory effects. However, due to the difference of the effects in group 1 and 2 was not that standing out, successively added silver ion to observe synergic effect of anti-inflammatory substances. Combined phenolic compound and silver ion showed dramatic effects in nitric oxide (NO) assay for increasing anti-inflammatory effects compared to other examined groups.

## 1. Introduction

Natural products are in the spotlight as they contain substances with anticancer and anti-inflammatory and antiallergic beneficial effects. Hetland et al. (2020) As natural products are originated from unprocessed natural extracts, and they show potential health benefits, eco-friendly properties, resulting in positive environmental and social effects than the substances created through chemical synthesis. In addition, the safety and effectiveness of using natural products have been proven through many clinical reports reported by different researchers. Gyawali et al. (2014)

Peanuts contain bioactive compounds with antioxidant activity, such as resveratrol, flavonoids, phenolic compounds, and phytosterols. Resveratrol (RE) is important and most researched natural polyphenols. RE has been described to exhibit a plethora of remedial benefits, including cardioprotective, antioxidant, anti-inflammatory, anti-hyperlipidemic, immuno-modulator, anti-platelet, vasorelaxant, and neuroprotective effects. Peanuts contain resveratrol and triterpene, and they exhibit anticancer, antioxidant activities. Candela (2020) RE and other compounds including flavonoids and phenolics has been widely used for various therapeutic benefits.

Silver Ions have been known for their various therapeutics’ effects for long. They have antibacterial, anti-inflammatory, and other various activities. Nanotechnology has provided many possibilities for preparing different forms of silver nanoparticles. The silver ions have high possibility for interactions with the cells. Therefore, keeping in mind the therapeutic effects of peanut’s leaf extract and silver ions, I conducted this study to investigate the synergistic effect of these both. Moreover, to find an effective method for cultivating peanuts containing high concentration of phenolic compounds, two groups of peanut plants were used in this experiment (with wound and without wound).

## 2. Materials and Methods

### 2.1 Peanut culture and extraction

The peanuts were cultivated at 28° C for 2 weeks following the controlled photoperiod (8h dark, 16h light). The control samples were grown at the same above-mentioned conditions without cutting or inoculation. The leaves of sample 2 were cut about 0.5cm with scissors to make a wound. Subsequently after wound, the culture was kept for two days with the same conditions and the leaves were removed. The leaves were treated with 0.1mg/ml ethanol, followed by shaking at 28° C for 12 hours. After 12 hours, the centrifugation was done to get the supernatant. The supernatant was concentrated using a rotary concentrator.

### 2.2 Analyzing total phenolic compound

The obtained concentrated extract (1 mg/mL) was diluted by adding 2 mL distilled water and mixed thoroughly with 0.5 mL of Folin–Ciocalteu reagent for 3 minutes. In the next step, 2 mL of 20% (w/v) sodium carbonate was added and the mixture was kept in dark 60 minutes and was used to measure the absorbance at 750 nm. The total phenolic content was calculated from the calibration curve and results were expressed in mg/g with fresh weight.

### 2.3 Analyzing total flavonoid

The total flavonoid content of the extract was determined through the aluminum chloride colorimetric method. Briefly 200 μL raw extract (1 mg/mL ethanol) was used with addition of 1 mL methanol, subsequently mixed with 4 mL of distilled water. NaNO2(0.3 mL, 5%) was added with AlCl3 solution (0.3 mL, 10%) in the mixture for 5 minutes following incubation. After incubation, the mixture was kept for 6 minutes and 2 mL of NaOH solution (1 mol/L ) was added. The final volume of the mixture was brought to 10 mL with double-distilled water. After this process, the mixture was kept for 15 minutes at room temperature, and absorbance was measured at 510 nm. The total flavonoid content was calculated using a calibration curve and the results were expressed as mg/g with weight.

### 2.4 DPPH assay

Scavenging activity of Hypericum extracts against DPPH radical was assessed according to the method of Blois with some modifications. Briefly, 1 ml of peanut extracts (0.01 mg dw/ml) was mixed with 4 ml of 0.005 mg/ml DPPH methanol solution. The reaction mixture was vortexed thoroughly and kept in the dark at room temperature for 30 minutes. The absorbance of the mixture was measured at 517 nm. Ascorbic acid and Butylated hydroxytoluene (BHT)were used as references. The ability to scavenge DPPH radical was calculated by the DPPH radical scavenging activity equation with the condition that control is the absorbance of DPPH radical in methanol and sample is the absorbance of DPPH radical solution mixed with sample extract/standard. All experiments were conducted in triplicates.

### 2.5 Cell culture

The macrophages (RAW 264.7) were purchased from the Korea Cell Line Bank (KCLB, Seoul, Korea) and were used in the experiment. Using Dulbecco Modified Eagle Medium (DMEM medium) containing 10% FBS and 1% P/S, the cells were cultured at 37° C with 5% CO2 conditions.

### 2.6 MTT assay

The cytotoxicity of peanut extracts 1 and 2 were measured using MTT assay. Cells were aliquoted into 96 well plates at 5×10^3^cells/well, and peanut extracts 1 and 2 were treated at 50 and 100 μg/mL, and silver at 20 μg/ml. After incubation for 24 hours, 20 μL of MTT [3-(4,5-Dimethylthiazol-2-yl)-2,5-diphenyl tetrazolium bromide] reagent (5 mg/ml) was added and incubated for 1 hour. After incubation, the supernatant was removed, and 100 μL of DMSO (dimethyl sulfoxide) was added to dissolve the formazan crystals. The absorbance was measured at 550 nm using a microplate reader. The cytotoxicity was expressed as a percentage calculated by the formula below compared to the untreated control group.

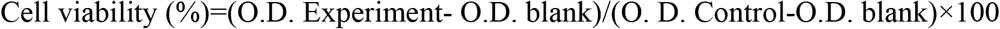

### 2.7 Nitirc Oxide (NO) assay

The amount of NO generated from RAW 264.7 cells were measured using Griess reagent [1% (w/v), sulfanilamide in 5% (v/v), phosphoric acid, and 0.1% (w/v) naphtylethylenediamine-HCl] in the cell culture medium. It was measured in the form of NO2- cells were aliquoted in a 96-well plate at 5×10^3^ cells/well, peanut extracts 1 and 2 were at 50 and 100 μg/mL, and silver at 20μg/ml, alone or in combination, and pretreated for 1 hour, 1 μg /mL LPS was treated and incubated for 24 hours. After transferring 50 μL of the culture medium to another 96 well plate, 100 μL of Griess reagents were mixed and reacted at room temperature for 10 minutes to measure absorbance at 550 nm. NO content was determined by the concentration of NO in the culture medium through a standard curve for each concentration based on sodium nitrite (NaNO2).

### 2.8 TNF-α ELISA analysis

To measure the content of TNF-a in the cell culture medium, BD OptEIA Mouse TNF-α ELISA kit (BDbioscience) was used. Peanut extract and silver were treated alone or in combination, and the cell culture medium of the control group and the provided standard sample were dispensed into each well and analyzed according to the method suggested by the manufacturer.

## 3. Results and Discussion

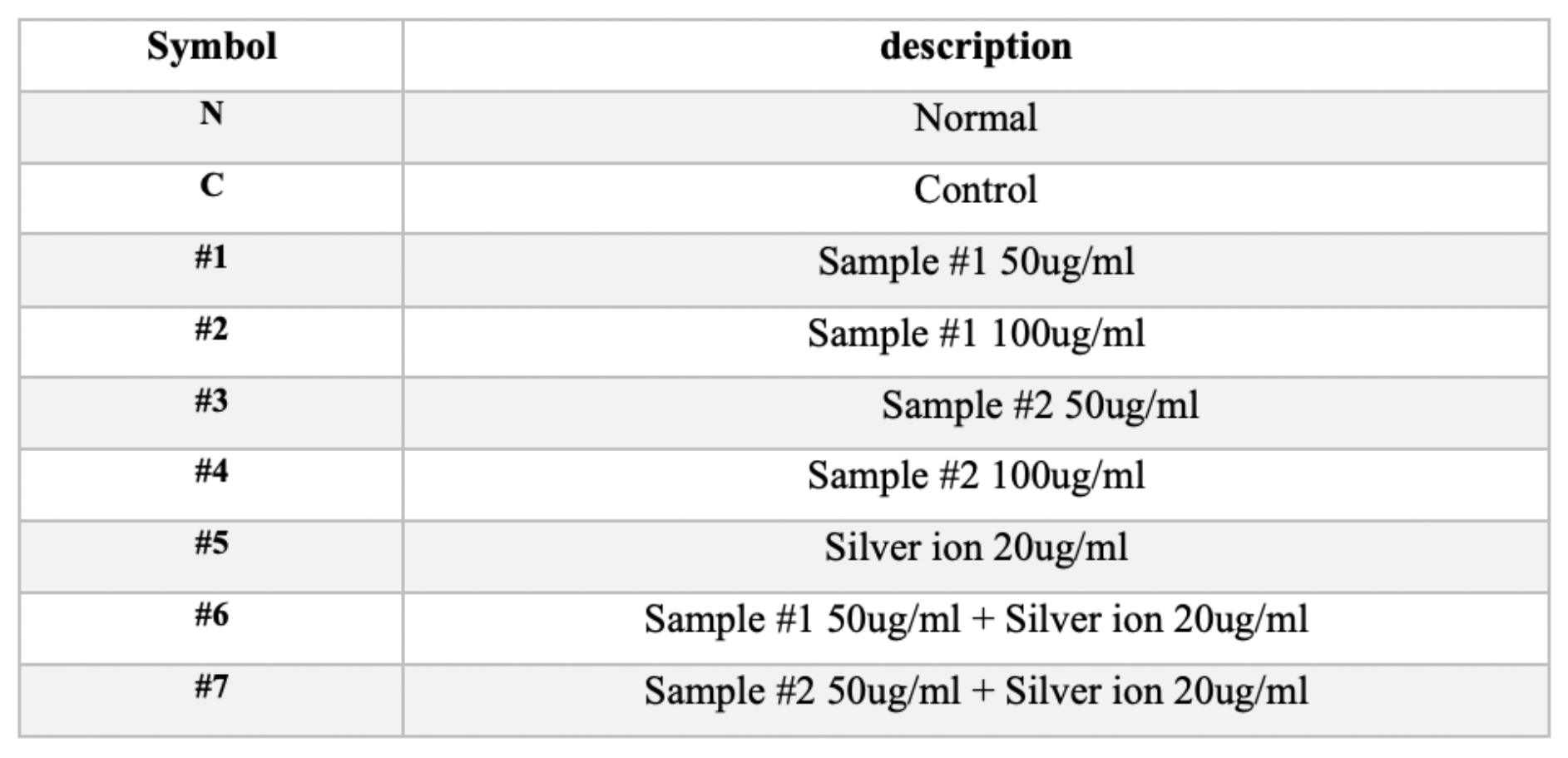

The above symbols are indicated in the figures below.

**figure 1.**
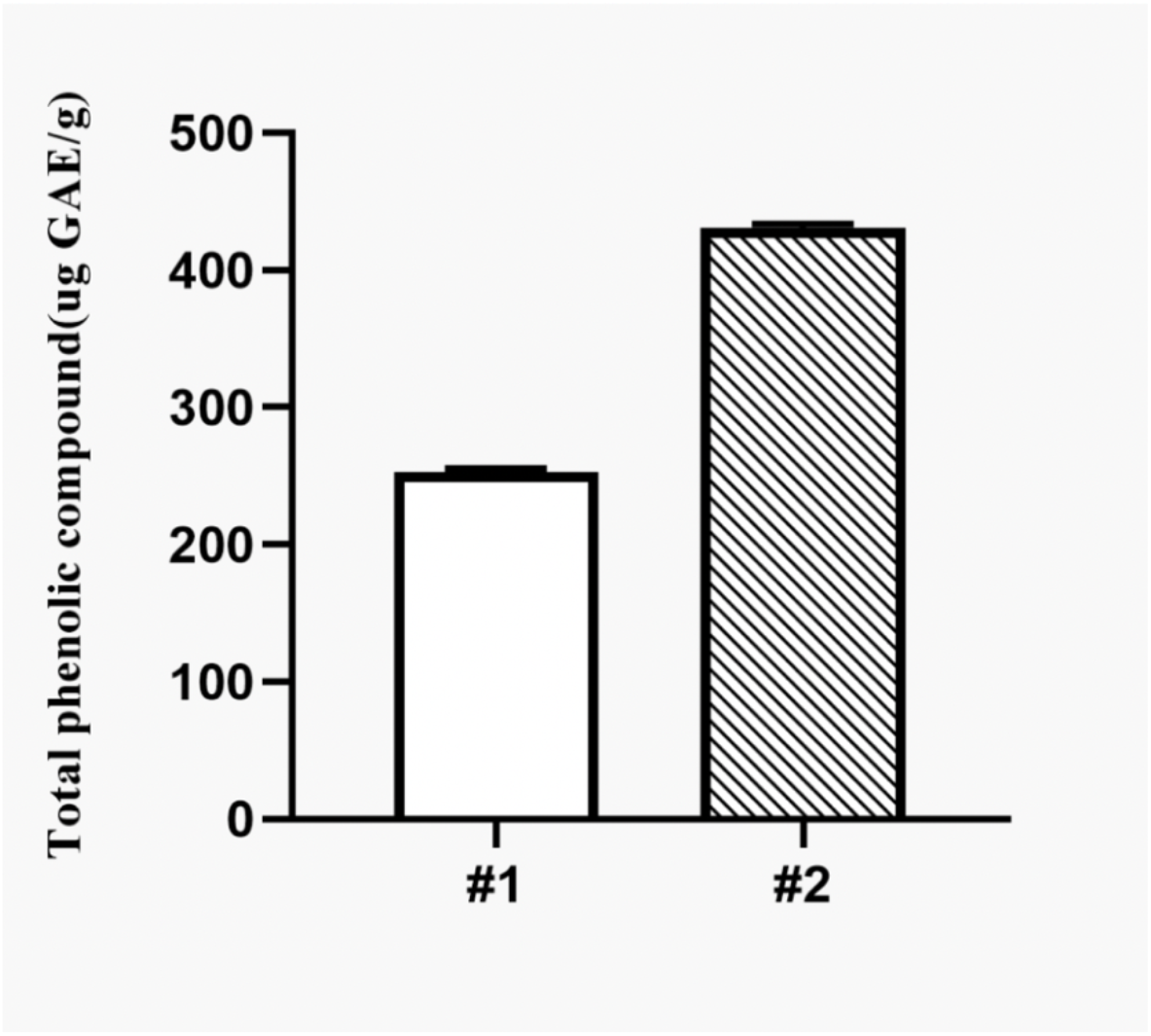
Total phenolic compound measurement

I conducted experiments on two groups of plants, group 1 without wounds and group 2 with wounds. The total phenolic content exhibited in group 2 was higher as compared to the group 1 plants. The plants with wound showed more amount of phenolic compound. Similarly, when I conducted and analyzed the antioxidant activity of both groups, group 2 showed higher antioxidant activity as compared to group 1.

**figure 2.**
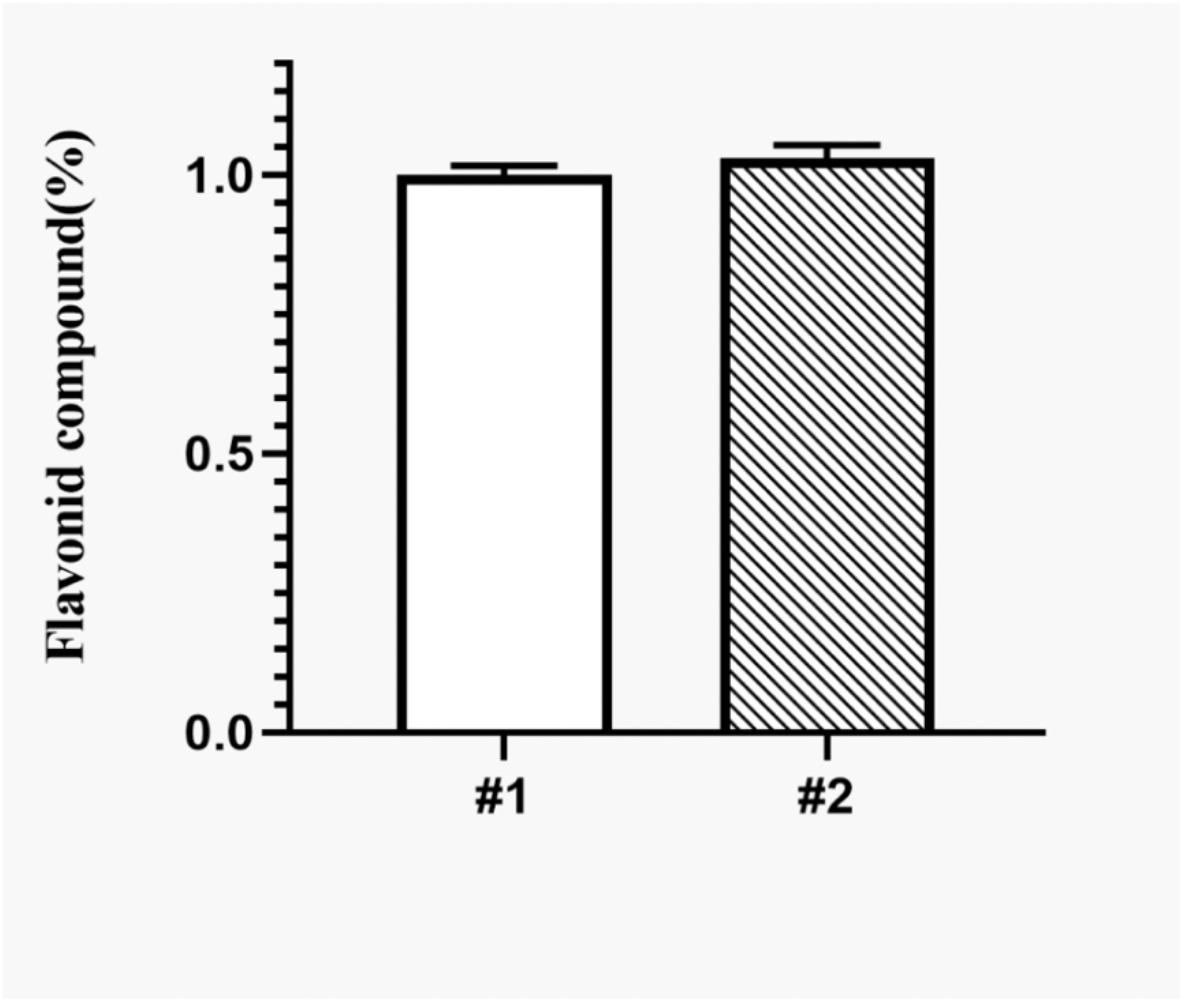
Flavonoid compound measurement

In the same trend, the total flavonoid contents were also produced in relatively large amounts in group 2. Accordingly, it has been reported that the plants with higher content of flavonoids exhibit relatively higher antibacterial, anti-inflammatory, and antihistamine activities.

**figure 3.**
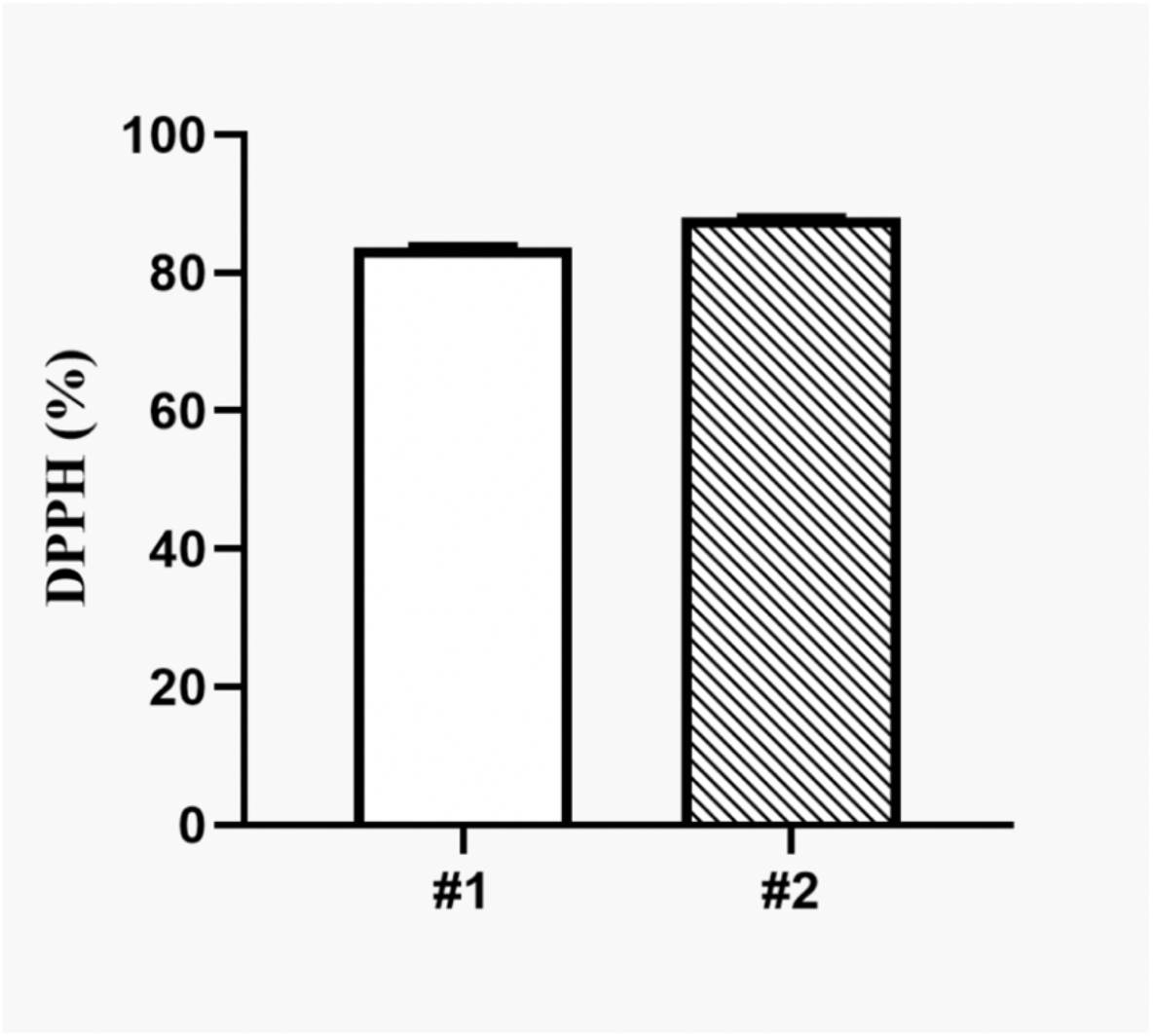
DPPH assay for antioxidant activity

DPPH free radical scavenging is an accepted mechanism for screening the antioxidant activity of plant extracts. According to my results, the wounded group of peanut plant extracts showed higher scavenging ability as compared to the group 1.

**figure 4.**
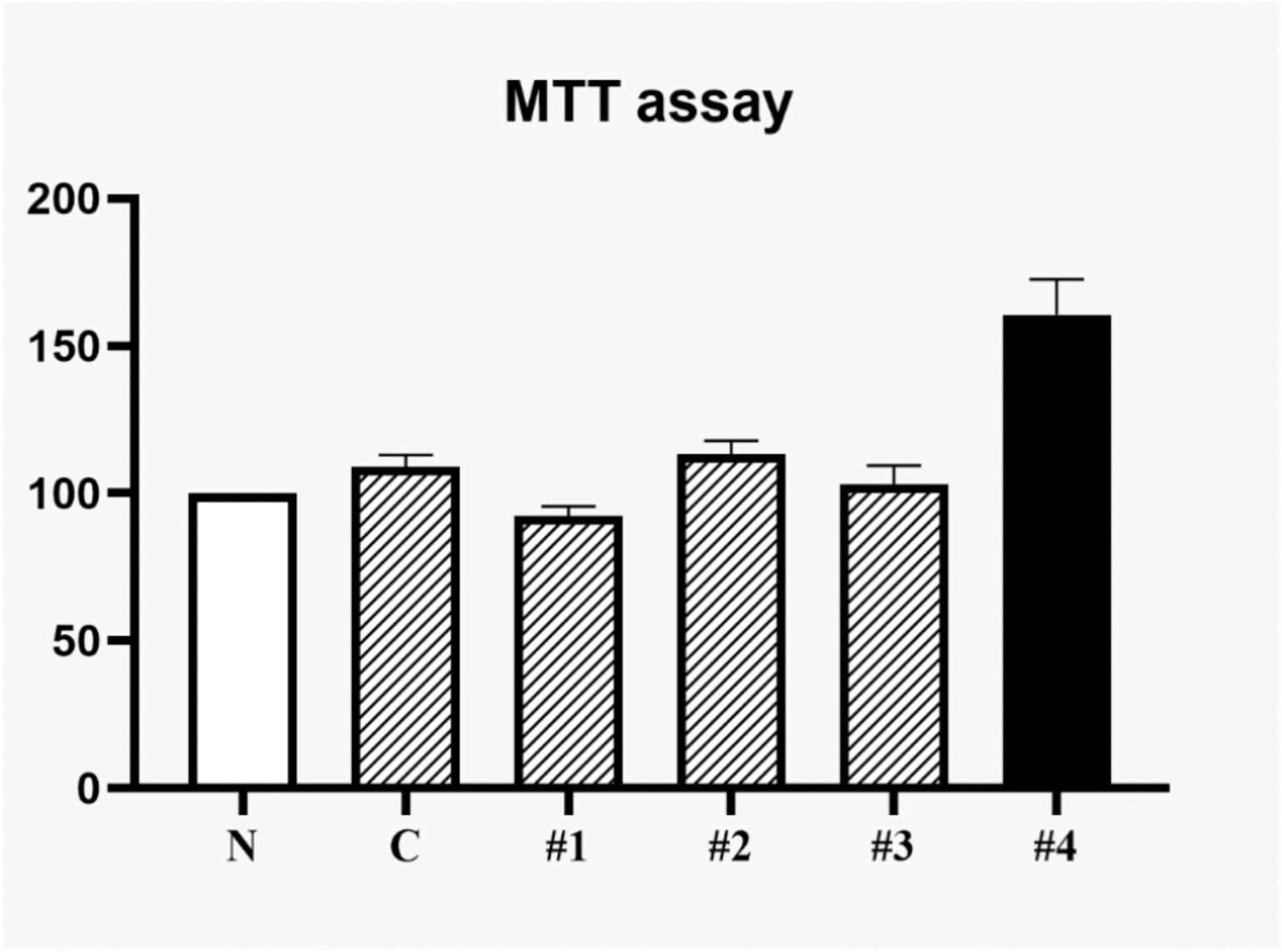
MTT assay

I used the MTT assay to measure cellular metabolic activity as an indicator of cell viability and cytotoxicity. The RAW 264.7 cells were used for MTT assay. In this study, Sample #2 100ug/ml, showed the highest cell survivability, and there is no significant difference between Sample #1 100ug/ml (#2) and Sample #2 50ug/ml (#3). It is interpreted as a number. Nevertheless, it can be noted that when comparing #3 and #1 that group 2 shows a higher level at the same 50ug/ml. However, there was no significant difference between control and #1, #2, #3, therefore, Sample #2 100ug/ml showed the most significant results. This indicates that the highest cell survivability was measured in #4.

**figure 5.**
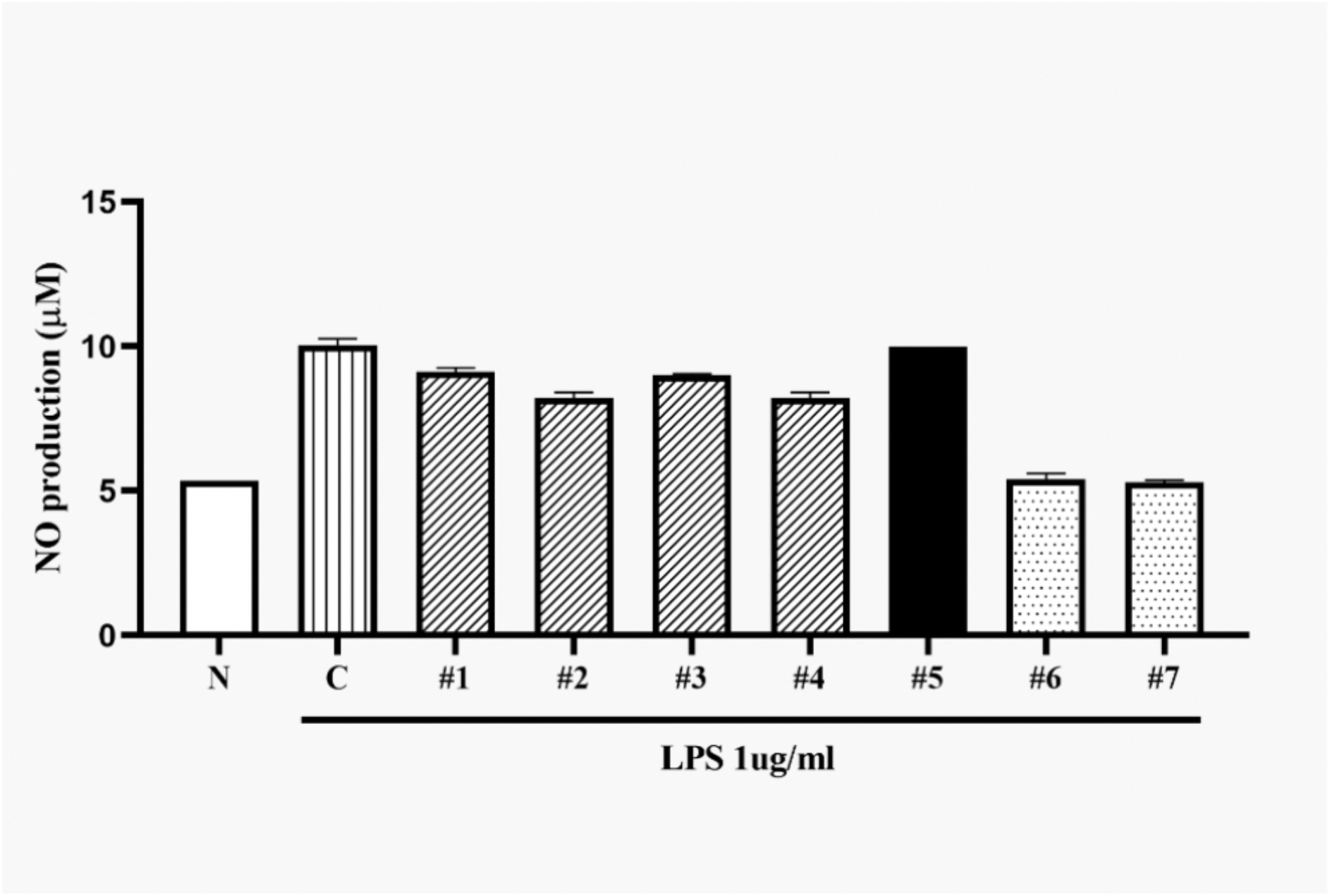
Nitric Oxide (NO) assay

It is well recognized that inflammation plays important roles in initiation and progress of many diseases including cancers in multiple organ sites. Controlling inflammation is one of the important strategies for the prevention and treatment of cancer. Higher levels of NO cause inflammation. I used NO assay for determination of nitrite and nitrate in plant extracts. I observed that the absorbance levels between control and #5 were similar. Moreover, #1 through #4 showed lower NO levels than control with similar degree.

However, in case of #6 (composed of Sample #1 50ug/ml + Silver ion 20ug/ml) and #7 (Sample #2 50ug/ml + Silver ion 20ug/ml), the results obtained were more significant and they showed lower NO levels. It is reported that the anti-inflammatory effect was higher due to the synergistic effect with the silver ion through the generation of about half of the NO of the control.

**figure 6.**
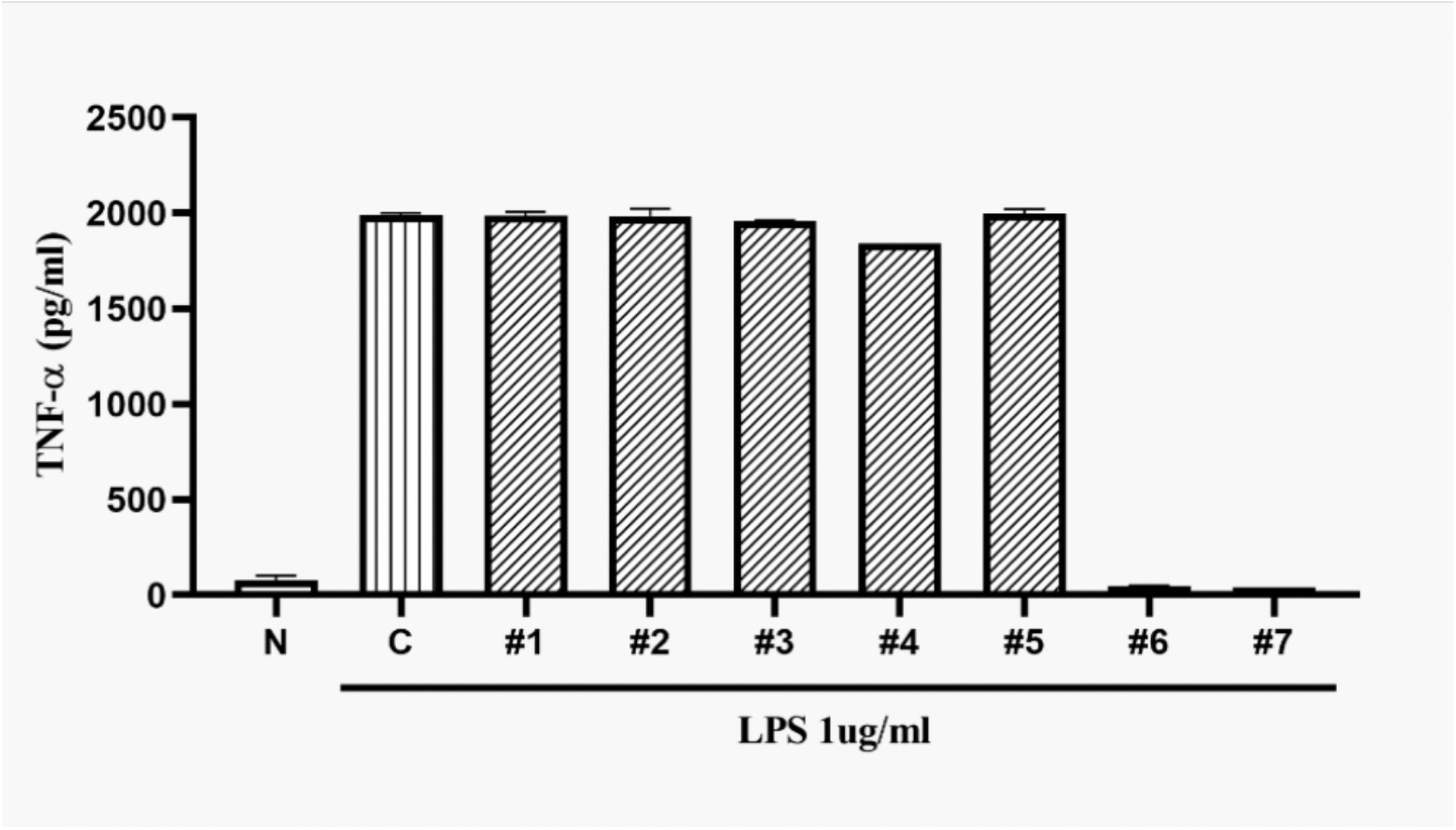
TNF-α ELISA analysis

It has been demonstrated in studies that pro-inflammatory cytokines such as TNF-α play vital roles in the progression of different diseases. Therefore, using ELISA assay I determined the effects of peanut extracts and their combination with silver ion. My results showed that the #1 through #5, no significant effect was observed as compared to Control, however, Sample #1 (50ug/ml) in combination with silver ion (20ug/ml) that is #6 and Sample #2 50ug/ml + Silver ion 20ug/ml that is #7 showed promising results. The #6 and #7 showed a lower number in the presence of silver ion. In the case of #7, which has a higher content with the addition of silver ion, it can be observed that it is lower than #6, which can be interpreted as an effective and potential anti-inflammatory result.

In conclusion, I observed that the wounded group 2 has a relative high efficacy compared to the group 1. This is because of plants that are undergoing the onslaught of wound-causing agents activate mechanisms of wound signaling and plants produces more compounds for healing and further defense. In addition, when #6 and #7 were treated in combination with silver ion, they showed more effectiveness. In this study, to determine whether the enhanced inhibitory effects from the combination group 1, 2 and silver ion were synergistic, I analyzed different assays. My results clearly demonstrated that the combination of extracts of group 1, 2 with silver ion produced synergistic effects in. TNF-α ELISA assay, No assay, and MTT assay. Jain et al. (2017). According to previous studies, too high flavonoid content may cause side effects. Alcalde-Eon et al. (2019) Accordingly, further studies are needed on flavonoid synthesis and efficacy at appropriate doses. The data from this study can be used to produce peanuts with a high polyphenol content, which can help advance medical technology such as cancer treatment.

